# A Positive Regulatory Feedback Loop Between EKLF/ KLF1 and TAL1/SCL Sustaining the Erythropoiesis

**DOI:** 10.1101/2021.04.08.439008

**Authors:** Chun-Hao Hung, Yu-Szu Huang, Tung-Liang Lee, Kang-Chung Yang, Yu-Chiau Shyu, Shau-Ching Wen, Mu-Jie Lu, Shinsheng Yuan, Che-Kun James Shen

## Abstract

The erythroid Krüppel-like factor EKLF/KLF1 is a hematopoietic transcription factor binding to CACCC DNA motif and participating in the regulation of erythroid differentiation. With combined use of microarray-based gene expression profiling and promoter-based ChIP-chip assay of E14.5 fetal liver cells from wild type (WT) and EKLF-knockout (*Eklf*^−/−^) mouse embryos, we have identified the pathways and direct target genes activated or repressed by EKLF. This genome-wide study together with molecular/cellular analysis of mouse erythroleukemic cells (MEL) indicate that among the downstream direct target genes of EKLF is *Tal1/Scl*. *Tal1*/*Scl* encodes another DNA-binding hematopoietic transcription factor TAL1/SCL known to be an *Eklf* activator and essential for definitive erythroid differentiation. Further identification of the authentic *Tall* gene promoter in combination with *<u>in vivo</u>* genomic footprinting approach and DNA reporter assay demonstrate that EKLF activates *Tall* gene through binding to a specific CACCC motif located in its promoter. These data establish the existence of a previously unknow positive regulatory feedback loop between two DNA-binding hematopoietic transcription factors that sustains the mammalian erythropoiesis.

## INTRODUCTION

Erythropoiesis is a dynamic process sustained throughout whole lifetime of vertebrates for the generation of red blood cells from pluripotent hematopoietic stem cell (HSC). In the ontogeny of mouse erythropoiesis, the major locations of HSC change orderly for several times, convert from embryonic yolk sac to fetal liver and then to the spleen and bone marrow in adult mice (1). At each of these tissues, the multistep differentiation process of erythropoiesis begins at the level of pluripotent hematopoietic stem cells (HSCs) and terminates with the production of erythrocytes (RBCs) and it is accompanied with a series of lineage-specific activation and restriction of gene expression. This stage-specific gene regulation cascade is mediated by several erythroid-specific/erythroid-enriched transcription factors, including GATA1, TAL1/SCL, NF-E2, and EKLF (2–5).

Among the factors regulating erythropoiesis lies the Erythroid Krüppel-like factor (EKLF/KLF1). EKLF is a pivotal regulator that functions both in erythroid differentiation and in controlling the lineage fate decision by the bipotential megakaryocyte-erythroid progenitors (MEPs) (6–10). *Eklf* is the first identified member of the KLF family of genes expressed in the erythroid cells, mast cells and their precursors (11, 12) as well as some of the other types of the hematopoietic cells but at low levels (6,13; Bio GPS). The critical function of *Eklf* in erythropoiesis has been demonstrated initially by gene abolition studies, with the *Eklf*-knockout mice (*Eklf*^−/−^) displaying severe anemia and died in utero at around embryonic (E) day 14.5 (E14.5) (14, 15). In addition to the dramatic decrease of the adult β globin gene expression, the molecular and cellular basis of lethality of *Eklf*^−/−^ mice is far more complex (8, 16). In particular, the impairment of the definitive erythropoietic differentiation is a major cause of embryonic lethality in *Eklf*^−/−^ mice in addition to β-thalassemia (17, 18). Significant reduction of the number of macrophages and abnormal macrophage morphology have also been observed in the E14.5 fetal liver of *Eklf*^−/−^ mice (13).

EKLF regulates its downstream genes, including the adult β globin genes, through binding of its C-terminal C2H2 zinc finger domain to the canonical binding sequence CCNCNCCC located in the promoters or enhancers (18–20) and the recruitment of co-activators, eg. CBP/p300 (21) and SWI/SNF-related chromatin remodeling complex (22, 23), or co-repressors, eg. mSin3A/HDAC1 (18) and Mi-2β/NuRD (7) complexes. Moreover, clinical associations exist between Eklf gene and different human hematopoietic phenotypes or diseases including β-thalassemia, the congenital dyserythropoietic anemia 4 (CDA4), neonatal anemia, the increased red blood cell protoporphyrin, hereditary persistence of fetal hemoglobin (HPFH), borderline HbA2, and inhibitor of Lutheran [In(Lu)] blood type (10, 24, 25). In erythroid progenitors, eg. CFU-E and Pro-E, EKLF is mainly located in the cytoplasm. Upon differentiation of Pro-E to Baso-E, EKLF is imported into the nucleus (20, 26) and form distinct nuclear bodies colocalized the β-globin locus, RNA polymerase II, SC35, and PML in discrete nuclear bodies (20, 27). In this way, EKLF participates in the spatial organization of chromatin configuration for efficient and coordinated transcription of genes including the β-globin locus in erythroid cells. More recently, genome-wide analysis of the global functions of mouse EKLF through identification of the direct transcription target genes has been conducted by using ChIP-Seq in combination with gene expression profiling (28, 29). The results from these studies suggest that EKLF functions mainly as a transcription activator in cooperation with TAL1/SCL and/or GATA1 to target genes including those required for terminal erythroid differentiation (28–30). However, much remains to be reconciled between the two studies with respect to the diversity of the genomic EKLF-binding locations and the deduced EKLF regulatory networks.

Besides EKLF, there are several other factors that have been shown to regulate erythropoiesis (2, 5). In particular, the T-cell Acute Lymphocytic Leukemia 1 (TAL1), also known as the Stem Cell Leukemia (SCL) protein, plays a central role in erythroid differentiation as well. The role of *Tal1/Scl* in primitive erythropoiesis has been demonstrated by the lethality of *Tal1*^−/−^ mice at E9.5 because of a complete absence of primitive erythrocyte in yolk sac (31, 32). Studies using erythroid cell lines (33) or adult-stage conditional *Tal1* gene knockout mice (34, 35) have shown the requirement of *Tal1* in definitive erythropoiesis. Another transcription factor known to play important roles in erythropoiesis is the zinc-finger DNA-binding protein GATA1, the consensus binding box ((T/A)GATA(A/G)) of which is present in the promoters and enhancer of most erythroid-specific genes (36–38). The cooperative functioning of TAL1 and GATA-1 in the regulation of erythroiepoiesis is closely associated with their physical associations at thousands of genomic loci (39).

Interestingly, *Eklf* appears to be a downstream target gene of the TAL1 factor. Whole-genome ChIP-seq analysis has identified the binding of TAL1 protein on the *Eklf* promoter in the primary fetal liver erythroid cells (40). Furthermore, there exists in the *Eklf* gene promoter the composite sequence of GATA-E box-GATA, which is a potential binding site of the GATA1-TAL1 protein complex required for the expression of *Eklf* gene in a transgenic mouse system (41). In the study reported below, we have combined promoter-based ChIP-chip technique using a high-specificity anti-EKLF antibody and microarray-based gene expression profiling to provide a genome-wide overview of the genes targeted by EKLF in the E14.5 mouse fetal liver cells. Remarkably, *Tal1* has turned out to be a direct target gene of EKLF, indicating the existence of a positive feedback loop between *Eklf* and *Tal1* for the regulation of erythropoiesis in mammals.

## MATERIALS AND METHODS

### Generation of Eklf^−/−^ mice

As described elsewhere (Hung et al., unpublished), the generation of B6 mouse lines with homozygous knockout of *Eklf* gene, *Eklf^−/−^*, was carried out in the Transgenic Core Facility (TCF) of IMB, Academia Sinica, following the standard protocols with use of BAC construct containing genetically engineered *Eklf* locus and E2A-Cre mice.

### Gene expression profiling by Affymetrix array hybridization

E14.5 mouse fetal livers from WT and *Eklf^−/−^* mouse fetuses were homogenized by repeated pipetting in phosphate-buffered saline (PBS) (10 mM phosphate, 0.15 M NaCl [pH 7.4]). Total RNAs were then isolated with Trizol reagent (Invitrogen) and subjected to genome-scale gene expression profiling using the Mouse Genome Array 430A 2.0 (Affymetrix, Inc.). Standard MAS5.0 method was applied to normalize the gene expression data. Gene expression values were log-transformed for later comparative analysis. Statistical analysis was carried out using R 3.0.2 language (R Development Core Team, 2013, http://www.R-project.org)

### Identification of differentially expressed genes

Genes with differential expression patterns between the WT and *Eklf*^−/−^ mice E14.5 fetal liver were first identified using two-sided two sample t-test with the significance level at 0.05. Since the set of probes for each annotated gene should all exhibit the same direction or sign when comparing the WT and *Eklf*^−/−^ samples, this consistency check was used to remove 257 ambiguous genes from the gene list. After filtering by the *p*-value threshold, a subset containing 12,277 statistically significant probe sets was obtained.

### Identification of EKLF-bound targets by using NimbleGen ChIP-chip array hybridization

The E14.5 mouse fetal liver cells were cross-linked, sheared, and the EKLF bound-chromatin complexes were immuno-precipitated (ChIP) with the AEK antibody (20) and rabbit IgG, respectively. DNAs were then purified from the immunoprecipitated chromatin samples by QIAquick PCR purification kit (Qiagen) and amplified by the Sigma GenomePlex WGA kit for hybridization with the Roche NimbleGen Mouse ChIP-chip 385K RefSeq promoter arrays.

There were 768,217 probes on the NimbleGen 385K ChIP-chip array. These probes were grouped into 21,536 sequence ids each of which contained 5 to 320 probes that ranged from 49 bp to 74 bp in length. The distances between the probes in the same sequence id ranged from 100 bp to 3,700 bp. In general, these sequence ids are located in the promoter regions of genes, roughly from −3.75kb to +0.75 kb relative to the transcription start site (TSS). The sequence id would be assigned a gene name when the gene’s coding sequence overlapped with the region from 10 kb upstream to 10 kb downstream of the sequence id. In this way, 652 sequence ids were found to be located in the intergenic regions and 20,884 sequence ids were near the coding regions of genes.

To identify the binding targets of EKLF, the moving window with size equal to 5 was adopted to test the hypothesis on positive mean value using one-sided t-test with the significant level at 0.0017. This smaller cut-off value was chosen to account for the multiple comparison. Specifically, there were 35 probes in each sequence id, and the adjusted *p*-value was derived by (1-(1-0.0017)^30^) ~0.05. The moving window was applied to each sequence id separately. Finally, the results were summarized at the sequence id level, and a sequence id would be defined as a target site of EKLF if there was a significant peak in the sequence id.

### Matching between Affymetrix probes and NimbleGen ChIP-chip probes

The E14.5 fetal livers gene expression data obtained from Affymetrix array hybridization analysis allowed us to further reduce the false positives from the ChIP-chip dataset. To do this, the annotation strategy used in annotating the sequence ids in ChIP-chip array was adopted to match the probes from these two platforms by gene symbols. After this procedure, there were a total of 78,634 matched pairs between the ChIP-chip sequence ids and Affymetrix probes.

### Co-occurrence of Binding Motifs and Relative Distance Distribution

226 known transcription factor-binding motifs were extracted from the previous report (28). For each of these binding motifs, the number of sequences in the mouse genome bearing the motif was sorted. The top co-existing binding motifs with EKLF were then further investigated. The relative distance between a co-existing motif and the EKLF-binding motif was calculated for each sequence id. However, when there existed multiple binding motifs in the same sequence id, multiple distances would be generated. In that case, the shortest distance was selected as the representative distance in that sequence id. The relative distance distribution was then plotted to inspect the potential localization biases. The existence of a localization bias provided further indication that two transcription factors might interact in certain way to regulate the particular target gene(s).

### Functional enrichment analysis

The analysis was carried out with use of IPA (Ingenuity^®^ Systems, www.ingenuity.com) to identify genes significantly associated with specific biological functions and/or diseases in the Ingenuity Knowledge Base. Right-tailed Fisher’s exact test was used to calculate the p-value determining the probability that each biological function and/or disease assigned to that data set was due to chance alone. The list of genes with significant EKLF-binding enrichment and deemed to be expressed differentially in WT and KO mice fetal livers was imported into IPA. The up-regulated and down-regulated EKLF targets were first mapped to the functional networks available in the IPA database, and then ranked by scores computed with the right-tailed Fisher’s exact test mentioned above. As listed in Supplemental Tables S3A and S3B, this analysis identified significant over-represented molecular and cellular functions (*p* value < 0.05) associated with the imported up-regulated and down-regulated EKLF targets that were eligible (score > 25) with the significance scores 26 and 34, respectively (Fig. S1 and S2).

### ChIP-qPCR

The ChIP-PCR analysis followed the procedures by Daftari *et al*. (42). The sonicated cell extracts from formaldehyde cross-linked E14.5 day mouse fetal liver cells were immuno-precipitated with anti-EKLF and purified rabbit IgG, respectiviely. The precipitated chromatin DNAs were purified and analyzed by quantitative PCR (qPCR) in the Roche LightCycle Nano real-time system. Sequences of the primers used for q-PCR designed by our lab are list in Supplementary Table 7. Each target gene was amplified with one set of primers flanking the putative EKLF-binding CACCC motif(s) and two sets of non-specific primers bracketing regions located at upstream and downstream of the CACCC motif(s), respectively.

### Plasmid construction

Mouse *Eklf* cDNA was derived by RT-PCR of RNA from DMSO-induced MEL cells and cloned into the vector pCMV-Flag (Invitrogen), resulting in pFlag-EKLF. Plasmids for luciferase reporter assay were constructed in the following way: *Tal1* promoter region from −1 to −900 relative to transcription start site of the newly identified *Tal1* exon 1 was amplified by PCR of mouse genomic DNA, with the addition of a XhoI cutting site at 5’ end and a HindIII site at 3’ end, and cloned into the XhoI and HindIII sites in the psiCHECK™-2 Vector (Promega) resulting in the plasmid p*Tal1*-Luc. *Tal1* promoter DNA fragments with the putative EKLF-binding CACCC box(es) mutated were generated by fusion PCR using the endogenous *Tal1* promoter as the template. Sequences of the three mutated CACCC boxes and their flanking regions in these fragments are: E1 box, 5’-CAGGCAAAACCAGGGACCAcatatTTAAAAATGATTCCCCTTCTCAAG-3’; E2 box, 5’-CAATAGCTCTTCAGTTAGCGGTGAAGGCTCATGAAcatatCCAC-3’; E3 boxes, 5’-GAGTTATTGACACAGCCCTGTcatatCCTCCCCCCACTG-3’. The inserts of all the plasmids were verified by DNA sequencing before use.

### Cell culture, differentiation, DNA transfection and knockdown of gene expression

Murine erythroleukemia cell line (MEL) was cultured in Dulbecco’s modified Eagle medium containing 20% fetal bovine serum (Gibco), 50 units/ml of penicillin, and 50 µg/ml of streptomycin (Invitrogen). For induction of differentiation, the cells at a density of 5×10^5^ /ml were supplemented with 2% dimethyl sulfoxide (DMSO; Merck) and the culturing was continued for another 24 to 72 hr. DNA transfection of the MEL cells and K562 cells was carried out using the TurboFect transfection reagent (Thermo Scientific) and Lipofectamine® 2000 transfection reagent (Life Technologies), respectively.

For knockdown of *Eklf* gene expression, MEL cell line-derived clones 4D7 and 2M12 (7) were maintained in 20 µg/mL of blasticidin (Invitrogen) and 1 mg/mL of G418 (Gibco). Differentiation of 4D7 and 2M12 cells was induced by 2% dimethyl sulfoxide (DMSO; Merck) for 48 hr. Expression of shRNA targeting and knocking-down *Eklf* was induced by the addition of 2 µg/ml of doxycycline (Clontech) for 96 hr, as described in Bouilloux et al. (7).

### RNA analysis

Total RNA from MEL cells and fetal liver suspension cells were extracted with TRIzol reagent (Invitrogen). cDNAs were synthesized using SuperScript II Reverse Transcriptase (RT) (Invitrogen) and oligo-dT primer (Invitrogen). Taq DNA polymerase was used for semi-quantitative RT-PCR analysis of the cDNAs. Quantitative real-time PCR (qPCR) analysis of the cDNAs was carried out using the LightCycler^®^ 480 SYBR Green I Master (Roche Life Science) and the products were detected by Roche LightCycler LC480 Real-Time PCR instrument. Primers used for qPCR analysis were designed following previous reports or from the online database PrimerBank: http://pga.mgh.harvard.edu/primerbank. Primers used for validating the microarray data and for *Tal1* exon 1 identification by RT-PCR were designed by our lab. The sequences of the DNA primers used in semi-quantitative RT-PCR and real-time RT-qPCR are available upon request.

### Western blotting analysis and antibodies

Whole-cell extract of MEL or mouse fetal liver cells were analyzed by polyacrylamide gel electrophories (PAGE) and Western blotting following the standard protocols. Enhanced chemiluminescence (ECL) detection system (Omics Biotechnology Co.) was used to visualize the hybridizing bands on blots. Goat anti-TAL1 antibodies, sc-12982 and sc-12984, were purchased from Santa Cruz, Inc. Anti-Flag (M2), anti-Tubulin(B-5-1-2), and anti-β-Actin (AC-15) mouse antibodies were purchased from Sigma-Aldrich. The anti-EKLF antibody (anti-AEK) was homemade (19).

### Reporter assay

For luciferase reporter assay in 293T cells, 1 µg of each of the wild-type p*Tal1*-Luc plasmid or its mutant forms were transfected into 4×10^5^ /ml of cells. The total amount of transfected DNA was kept at 0~3 µg with addition of 0~3 µg of empty vector pCMV-Flag. After 24 hr, the luciferase activities were measured using the Dual-Luciferase® Reporter Assay System (Promega). Firefly luciferase activity was used as an internal control and Renilla activity was used to monitor the transactivity of the *Tal1* promoter or its mutant forms.

### *In vivo* genomic footprinting

The status of nuclear factor-binding in the living MEL cells was investigated by dimethyl sulfate (DMS) cleavage *in vivo* and ligation-mediated PCR (LMPCR) as described previously (43,44) with some modification. The distal promoter region was analyzed with primer set D (P1[5’-885 GCTCACAAA CT CCT GTTTCAGAGGAG-860 3’], P2 [5’-867CAGAGGAGCTAATGTT CTGCCTTCTTC-840 3’], and P3 [5’-867 CAGAGGAGCTAATGTGTCTGCCTTCTTCTAG-837 3’]); the central promoter region was analyzed with primer set C (P1[5’-817GACATTAATACAGGCAAAACCAGG GACC-790 3’], P2[5’-798CCAGGGACCAC ACCCTTAAAAATGATTCC-770 3’], and P3[5’-798CCAGGGACCAC ACC CTTAAAAATGATTCCCC-768 3’]); the proximal promoter region was analyzed with the primer set P(P1[5’ 320GAAAGAAAAACCCAG A TACTCCTCAGC-294 3’], P2 [5’-287GGTTCCTACAATGTACCTATGG GCTTC-261 3’], and P3[5’-287GG TTCCTACAATGTACCTATGGGCTTCAATG-257 3’]). Different batches of DMS-treated cells were analyzed several times to check for consistency of the protection patterns. The relative intensities of the bands on the autoradiographs were estimated in a AlphaImager 2200 (Clontech).

## RESULTS

### Genome-wide identification of EKLF target genes by microarray hybridization and promoter-based ChIP-chip analyses

We first carried out gene profiling analysis to identify genes regulated by EKLF. The microarrays were hybridized with cDNAs derived from E14.5 fetal liver RNAs of four wild-type (WT) and four *Eklf* knockout (KO or *Eklf*^−/−^) embryos, respectively (Fig. 1A). Overall, there were 6,975 genes with differential expressions levels between the WT and KO mice (Fig. 1B, 1C, and 1D). Notably, the confidence index of the microarray hybridization analysis was approximately 70%, as the upregulation/downregulation of 8 out of 12 genes could be validated by semi-quantitation RT-PCR analysis (Fig. S3).

**Figure 1.**
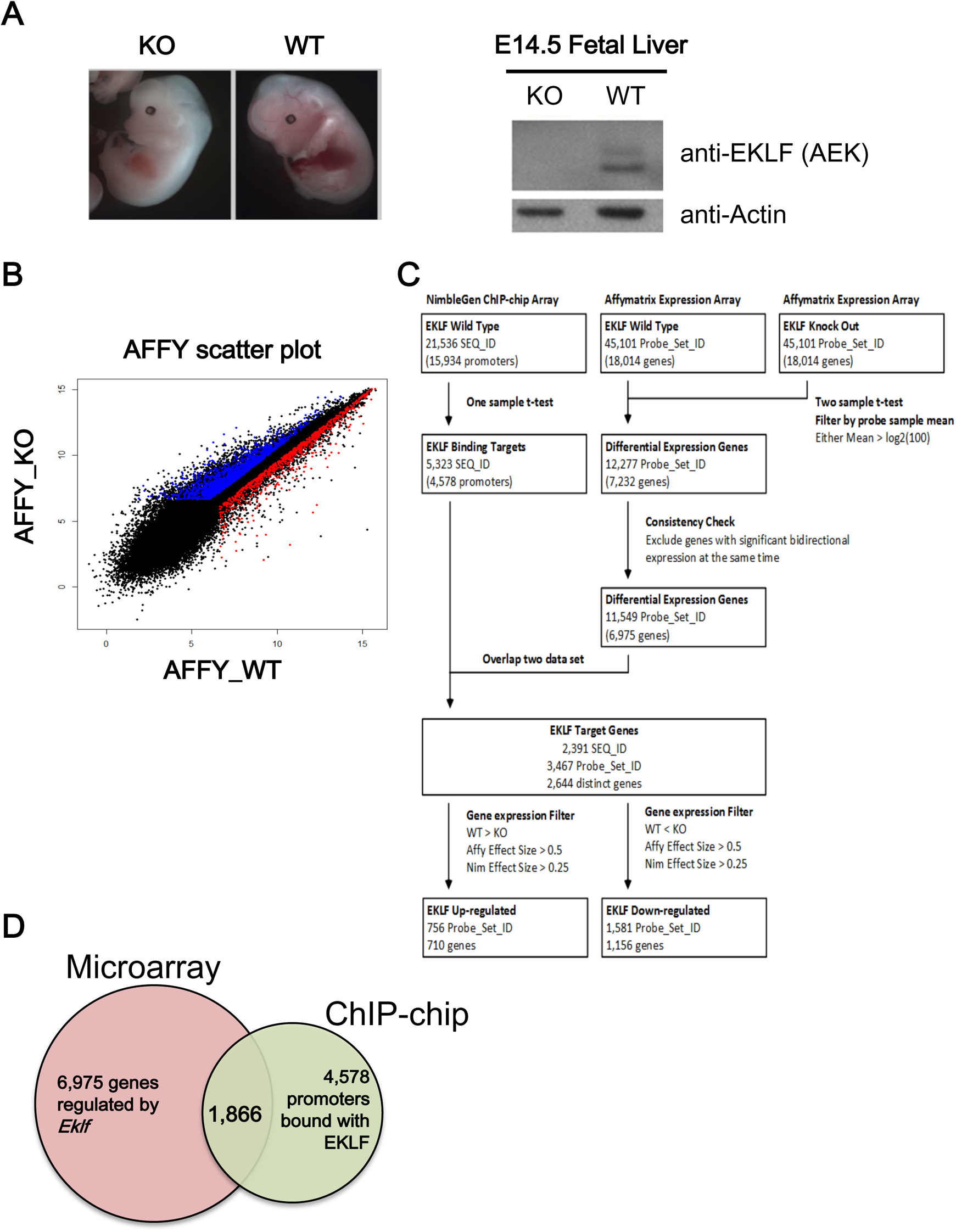
Identification of EKLF target genes by global gene expression profiling. **(A)** Left panels, representative appearance of E14.5 embryos of wild-type and *Eklf*^−/−^ mice. Right panels, Western blotting patterns of EKLF protein in E14.5 fetal lives. Actin was used as the gel loading control. **(B)** Scatter plot comparing the gene expression profiles of E14.5 fetal liver cells of WT and *Eklf*^−/−^ mice by Affymatrix array hybridization. Each gene on the arrays is displayed as a single dot on a logarithmic (log2) graph. The genes up-regulated and down-regulated by EKLF are indicated by the red and blue dots, respectively. **(C)** Overview of the workflow of ChIP-chip and microarray expression profiling. The flow chart illustrates the procedures used for analysis of the NimbleGen promoter ChIP-chip data and Affymatrix differential expression profiling data. The number of genes (SEQ_ID or Probe_Set) after data processing at each step is indicated in parentheses. **(D)** The Venn diagram showing the overlapping between gene sets derived from the microarray hybridization analysis (6,975 genes) and ChIP-chip analysis (4,578 genes), respectively, of E14.5 fetal liver cells of WT and *Eklf*^−/−^ mice.

We then performed ChIP-chip analysis (Fig. 1C) using a promoter-based microarray and the high-specificity polyclonal anti-mouse EKLF antibody (anti-AEK) (19, 26); (Fig. 1). The probes on the ChIP-chip array were grouped into 21,536 sequence ids (SEQ_ID). The promoter of each annotated gene was defined as the region from −3.75kb upstream to 0.75 kb downstream of the transcription start site (TSS). The SEQ_IDs with at least one significant peak were defined as the potential target binding sites of EKLF. Overall, enriched EKLF-binding was present in 5,323 SEQ_IDs corresponding to 4,578 promoters (Fig. 1C and Fig. 1D). We also validated the ChIP-chip data by ChIP-qPCR. 9 of 13 promoters on the ChIP-chip list indeed were bound with EKLF as shown by this assay (Fig. S4).

The data from the ChIP-chip and the microarray gene profiling experiments were then combined to identify the putative EKLF target genes. After matching between the 11,549 differentially expressed probe sets from the microarray data and the 5,323 significant SEQ_IDs from the ChIP-chip array data, 2,391 SEQ_IDs (11.1%) from the ChIP-chip data and 3,467 probe sets (7.7%) from the microarray hybridization data remained. In the end, combination of the two data sets resulted in 2,644 distinct genes (Fig. 1C and Table S2A). This gene list included only genes with altered expression level in the *Eklf*^−/−^ fetal liver at the *p* < 0.05 level and with at least one statistically significant EKLF-binding site (*p* < 0.0017) in the promoter region without considering the fold change of expression and enrichment of binding.

Upon filtering with the effect sizes of the ChIP-chip data (>0.25) and microarray profiling data (> 0.5), the number of EKLF-bound and regulated targets was reduced from 2,644 to 1,866, among which 1,156 were down-regulated and 710 were up-regulated in E14.5 WT fetal liver (Fig. 1C, Fig. 1D, Table S2B and Table S2C). The above data supported the scenario that the promoter-bound EKLF could function as either repressors or activators *in vivo*. Notably, the promoters bound with and regulated by EKLF were distributed throughout the mouse genome with no obvious preference for any chromosome (Fig. 2A and 2B).

**Figure 2.**
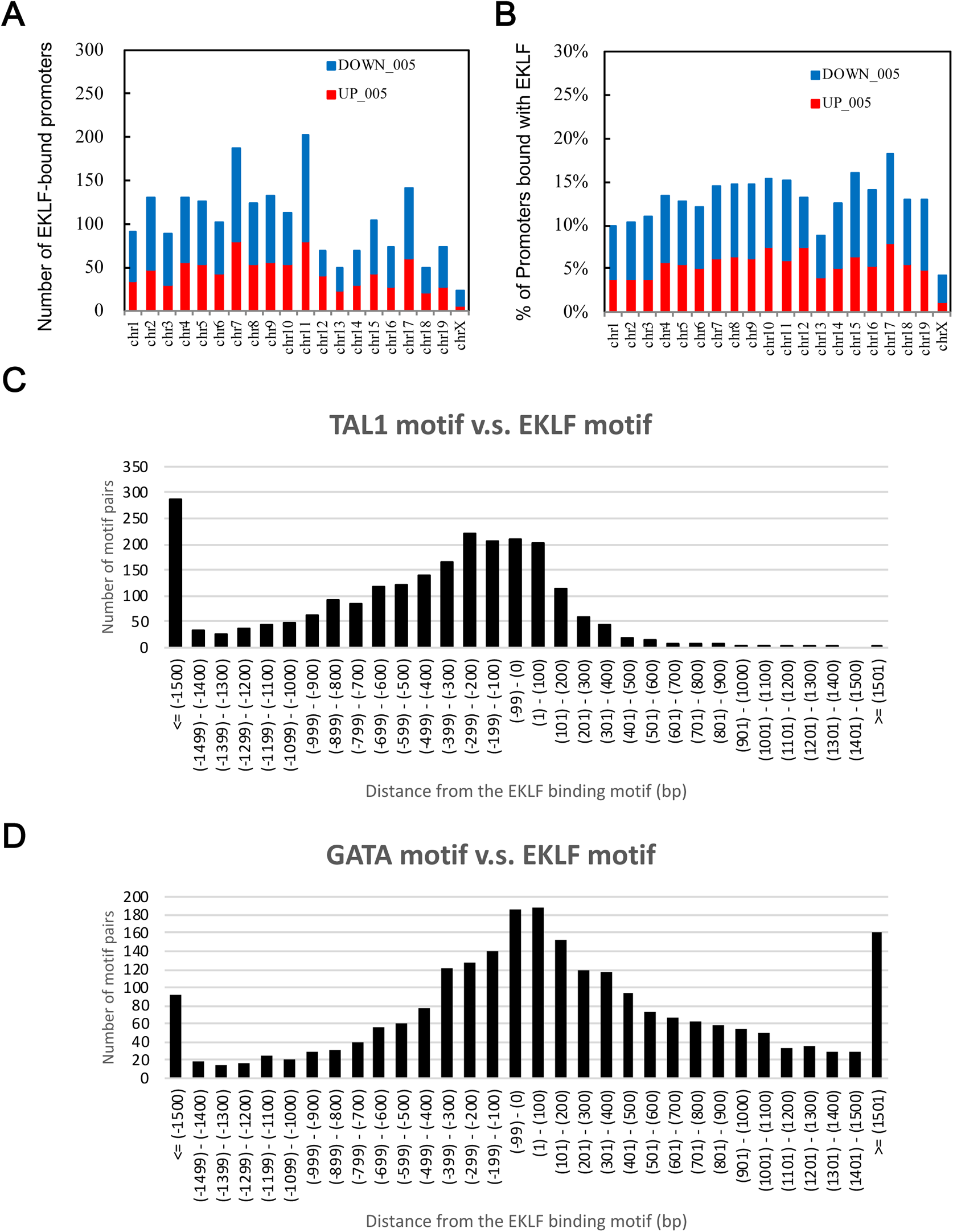
Chromosome distribution patterns from global analysis of direct target genes of EKLF in E14.5 fetal liver cells. **(A)** The numbers of the putative EKLF-bound promoters on the different mouse chromosomes. **(B)** The percentages of promoters of the individual mouse chromosomes bound with EKLF. **(C)** and **(D)** Distributions of the distances between the binding motif of TAL1 (C) or GATA1 (D) and that of EKLF on the mouse genome in E14.5 fetal liver cells. Upstream locations are indicated by the “-” sign, while the downstream locations are indicated by the “+” sign.

### Functional and pathway analysis of EKLF target genes

Previously reports by others using the Ingenuity Pathway analysis (IPA) and GeneGo MetaCore analysis platform showed that EKLF target genes were associated with a variety of cellular activities or pathways including general cellular metabolism, cell maintenance, cell cycle control, DNA replication, general cell development and development of hematologic system (29, 47). To gain further insight into the potential biological roles and functions of EKLF, we applied the IPA software for analysis of the putative 1,866 direct target genes of EKLF. The analysis identified the top five over-represented networks of the down-regulated EKLF targets and up-regulated EKLF targets, respectively (Table 1). The significance of the relevant networks were strengthened with use of the higher cut-off score of 25 to ensure that reliable functional networks built by IPA were eligible (Table 1, Supplementary Table S3A and S3B). This network analysis by us confirmed the previously established association of the hematological system development/function with the up-regulated EKLF targets (29). Notably, the top network associated with either the down-regulated EKLF targets or up-regulated EKLF targets was related to metabolism and small molecule biochemistry (Table 1). Additionally, the significant networks/functions associated with the down-regulated EKLF targets were more broad than those with the up-regulated EKLF targets (Table 1). Overall, our analysis was consistent with the previous studies (28, 29) in that the promoter occupancy by EKLF also played important roles in developmental processes, other than erythropeisis and hematological development.

We also used IPA software to group the number of EKLF targets according to their respective biological functions, regulatory pathways and physiological functions (Tables 2, S4A and S4B). Of the top five molecular and cellular functions, the cell death/survival, cellular assembly/organization, and cellular function/maintenance were overrepresented in both the up-regulated as well as down-regulated EKLF targets. Further enrichment analysis of the canonical pathways using the IPA software revealed significant overrepresented pathways across the same two genes lists (Tables 3, S5A and S5B). The prominent enrichment of the up-regulated EKLF targets were related to cell cycle control of chromosomal replication, EIF2 signaling, mitochondrial dysfunction, hypusine biosynthesis, and tryptophan degradation III (Eukaryotic) (Supplementary Fig. S5). The enrichment of the down-regulated EKLF targets was related to insulin receptor signaling, chondroitin sulfate degradation (Metazoa), gap junction signaling, nitric oxide signaling in the cardiovascular system, and PDGF signaling (Supplementary Fig. S6). The above together further established the specific functions and associated biological pathways associated with the EKLF target genes.

### Identification of potential transcription factors co-regulating the EKLF targets

Since co-occurrences of specific transcription factor-binding motifs in the promoters would suggest the cooperation of these factors in transcriptional regulation (48, 49), we searched factor-binding motifs across the EKLF-bound promoters as described in Material and Methods. Specifically, the consensus transcription factor-binding motifs were ranked based on how often a particular motif occurred within the sequence id. This observed frequency was applied to all the consensus transcription factor-binding motifs identified within the EKLF-bound regions on each sequence id (Table 4 and Supplementary Table S6). As expected, the most abundant transcription factor-binding motif in the EKLF-bound and regulated promoters was the consensus EKLF-binding sequence CACCC, which was present a total of 2,390 times in 2,391 of EKLF target sequence ids corresponding to 2,143 times in 2,644 distinct gene promoters. Consistent with Tallack et al. (28), the binding motifs of known transcription factors functionally interacting with EKLF, such as TAL1 and GATA1 (28), were also identified, which were present at least once in 2,390 (2,143 distinct gene promoters) and 2,384 (2,139 distinct gene promoters) of the EKLF target sequence ids, respectively. In addition, the binding motifs of a number of other transcription factors possibly interacting with EKLF functionally, such as PEA3, LVa, H4TF-1 and XREbf, etc., were also identified in this way (Tables 4 and S6).

To investigate the functional cooperation between TAL1 and EKLF or between GATA1 and EKLF, we further analyzed the distance between the binding motifs of GATA1 or TAL1 and that of EKLF. Indeed, the distribution of TAL1 binding motifs has the highest frequencies between +100 bp and −100 bp from EKLF binding motifs, indicating a functional cooperation between TAL1 (or possibly Ldb1 complex) and EKLF. Moreover, this cooperation likely acts through the binding of TAL1 at upstream of EKLF protein (Fig. 2C). The distribution pattern of GATA1 binding motifs also supported the cooperation between this factor and EKLF (Fig. 2D), although there is no obvious upstream/downstream preference between these two factors.

### Likelihood of *Tal1* gene as a regulatory target of EKLF in E14.5 fetal liver cells

Our motif analysis across the EKLF-bound promoters revealed that the binding motifs of the transcriptional factor TAL1 had the highest frequency of co-occupancy with the binding motifs of EKLF (Table 4), suggesting a functional cooperation between these two factors in transcriptional regulation. Interestingly, such cooperation in the hematopoietic system were often associated with the transcriptional activation of one partner by another partner (29, 30). For instance, either the EKLF-GATA1 or TAL1-GATA1 duet served as part of specific activation complex(es) in the erythroid cells (49–51), and both the *Eklf* gene (52, 53) and the *Tal1* gene (54) were activated by the GATA1 factor. We thus further investigated whether there was also an epistatic relationship between *Eklf* and *Tal1*.

Gene expression profiling by microarray hybridization revealed a 2.5-fold (effect size=1.3139) down-regulation of the *Tal1* transcript in E14.5 *Eklf^−/−^* fetal liver in comparison to the wild-type E14.5 fetal liver (Supplementary Table S2A). This microarray data was validated by RT-qPCR. As shown, the level of *Tal1* mRNA in the *Eklf^−/−^* fetal livers was decreased significantly, down to 47% of the level detected in wild-type fetal liver (left histograph, Fig. 3A). In parallel, the TAL1 protein level in the *Eklf^−/−^* fetal livers cells was also down-regulated, by 70%, when compared to the wild type (right panels and histograph, Fig. 3A). Thus, not only TAL1 factor could activate the *Eklf* gene transcription (40–41), but the *Tal1* gene might also be a regulatory target of EKLF. Since *Eklf^−/−^* mice and *Tal1*^−/−^ mice both exhibited a deficit of erythroid-lineage cells after the stage of basophilic erythroblasts (34, 47), we suspected that the promotion of erythroid terminal differentiation from pro-erythroblasts to basophilic erythroblasts very likely required EKLF-dependent activation of the *Tal1* gene transcription.

**Figure 3.**
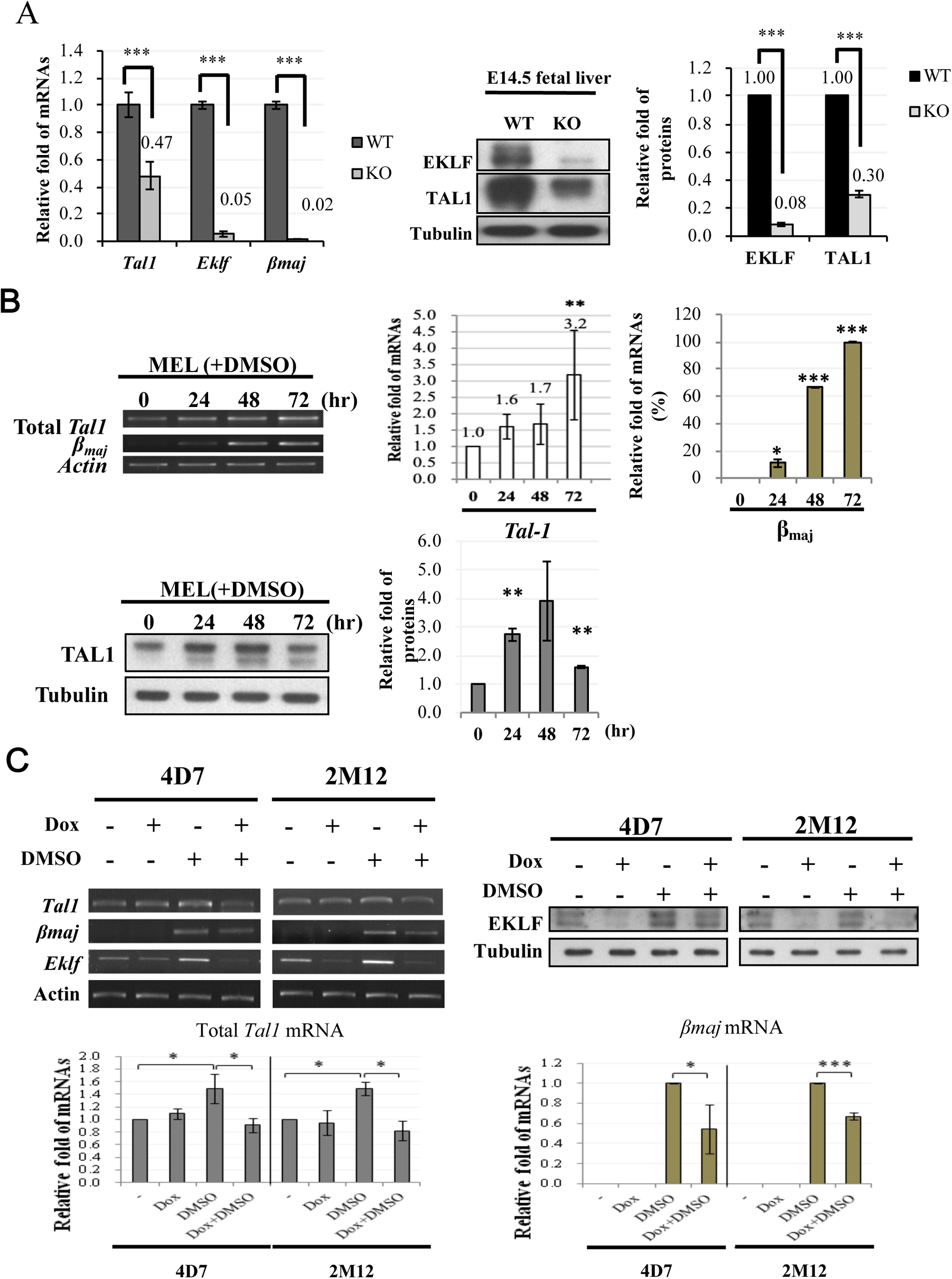
Tal1 as a direct target gene of EKLF. **(A)** Left, bar diagram of the relative mRNA levels of *Tal1*, *Eklf* and *βmaj* in E14.5 fetal liver cells of the WT and *Eklf*^−/−^ (KO) mice, as analyzed by RT-qPCR. *** p < 0.001 by t test. Error bars, S.E.M. Middle panels and right histobar diagram, Western blotting analysis of TAL1 and EKLF in E14.5 fetal livers of WT and *Eklf*^−/−^ (KO) mice. Tubulin was used as the loading control. *** p < 0.001 by t test. Error bars, STD. **(B)** Top, expression levels of *Tal1* and *βmaj* in MEL cells without or with DMSO induction for 72 hr. The gel patterns of the semi-quantitative RT-PCR bands are shown on the left, and the histographs of the statistical analysis of the data are shown on the right. * p< 0.05, ** p < 0.01, and ***p<0.001 by t test. Error bars, S.D. Bottom, Western blotting analysis of the levels of TAL1 protein in MEL cells during DMSO-induced differentiation. Tubulin was used as the loading control. The statistical analysis of the data is shown in the bar diagrams on the right. ** p < 0.01 by t test. Error bars, S.D. **(C)** Analysis of gene expression in 4D7 and 2M12 cells without and with doxycycline (Dox)-induced expression of *Eklf* shRNA. The cells without or with induction by DMSO for 48 hr were treated with doxycycline. The levels of EKLF protein in the whole cell extracts were then analyzed by Western blotting, as exemplified in the upper right panels. Tubulin was used as the loading control. RT-PCR analysis showed that knockdown of EKLF reduced the levels of total *Tal1* mRNA and *βmaj* mRNA in DMSO-induced cells, as exemplified in the upper left panels and statistically analyzed in the two histobar diagrams below. The gel band signals were all normalized to that of actin. * p< 0.05, ** p < 0.01, and ***p<0.001 by student t test. Error bars, S.D.

### EKLF as an activator of Tal1 gene expression during erythroid differentiation

To further examine whether EKLF was an activator of *Tal1* gene transcription in erythroid cells, we first analyzed the expression level of *Tal1* mRNA in cultured mouse erythroid leukemic (MEL) cells during DMSO induced erythroid differentiation. Similar to βmaj mRNA, the *Tal1* mRNA was expressed in un-induced MEL at a basal level, which was increased by 2-3 folds upon DMSO differentiation (top, Fig. 3B). In consistency, the protein level of TAL1 was also up-regulated during an 48 hr period of DMSO-induced differentiation, but down-regulated subsequently (bottom, Fig. 3B). The up-regulation of the *Tal1* gene supported the scenario that sustained higher expression of the *Tal1* gene was required for erythroid differentiation. The biphasic expression profile of the TAL1 protein further suggested that the requirement of TAL1 for MEL cell differentiation was up to 48 hr after DMSO-induction, which corresponded to the basophilic / polychromatic stages of erythroid differentiation.

We then analyzed *Tal1* mRNA levels in two independent MEL cell-derived stable clones, 4D7 and 2M12. As shown in Fig. 3C, knock-down of *Eklf* mRNA by the doxycycline-induced shRNAs led to a significant reduction of the *Tal1* mRNA under the condition of DMSO-induced erythroid differentiation, but not in un-induced MEL cells. The latter result further supported that EKLF was not part of the regulatory program of *Tal1* gene transcription in MEL cells prior to their differentiation. The data of Fig. 3C indicated that EKLF was required for the activation of *Tal1* gene transcription during DMSO-induced erythroid differentiation of MEL cells. Together with the loss-of-function of *Eklf* study in mouse fetal liver (Fig. 3A), we conclude that while TAL1 is a known activator of *Eklf* gene transcription, EKLF also positively regulates *Tal1* gene transcription during erythroid differentiation from CFU-E/pro-erythroblasts to the basophilic / polychromatic erythroid cells.

### Binding *in vivo* of EKLF to the upstream promoter of *Tal1* Gene

How would EKLF activate the *Tal1* gene transcription during erythroid differentiation? It could either directly activate the *Tal1* gene through DNA-binding in the regulatory regions of the gene, eg. its promoter or enhancer, or indirectly through other transcriptional cascades. In interesting association with the above data of *Tal1* expression in the presence and absence of EKLF, the ChIP-chip analysis identified two regions with significant reads of EKLF-binding, one of which (region I,114,551,700 - 114,552,900 on chromosome 4, NCBI 36/mm8) was located around the *Tal1* gene in E14.5 fetal liver cells (Fig. 4A, Table S1, Table S2A). In mouse erythroid cells, the *Tal1* gene encodes a *Tal1* mRNA isoformA (Fig. 4B) consisting of 5 exons, with the most upstream exon1 located at 115,056,426 - 115,056,469 (NCBI 36/mm8). However, no CCAAT box or TATA box or CACCC box could be found within 300 bp upstream of this exon 1. Instead, we found these motifs in a region ~ 860 bp upstream of isoform A exon 1 (Fig. 4B and 4C; see also sequence in Fig. 5A). We thus suspected that exon 1 of *Tal1* gene might be longer than currently documented in the database Alternatively, there might be another exon upstream of the exon 1 of isoform A.

**Figure 4.**
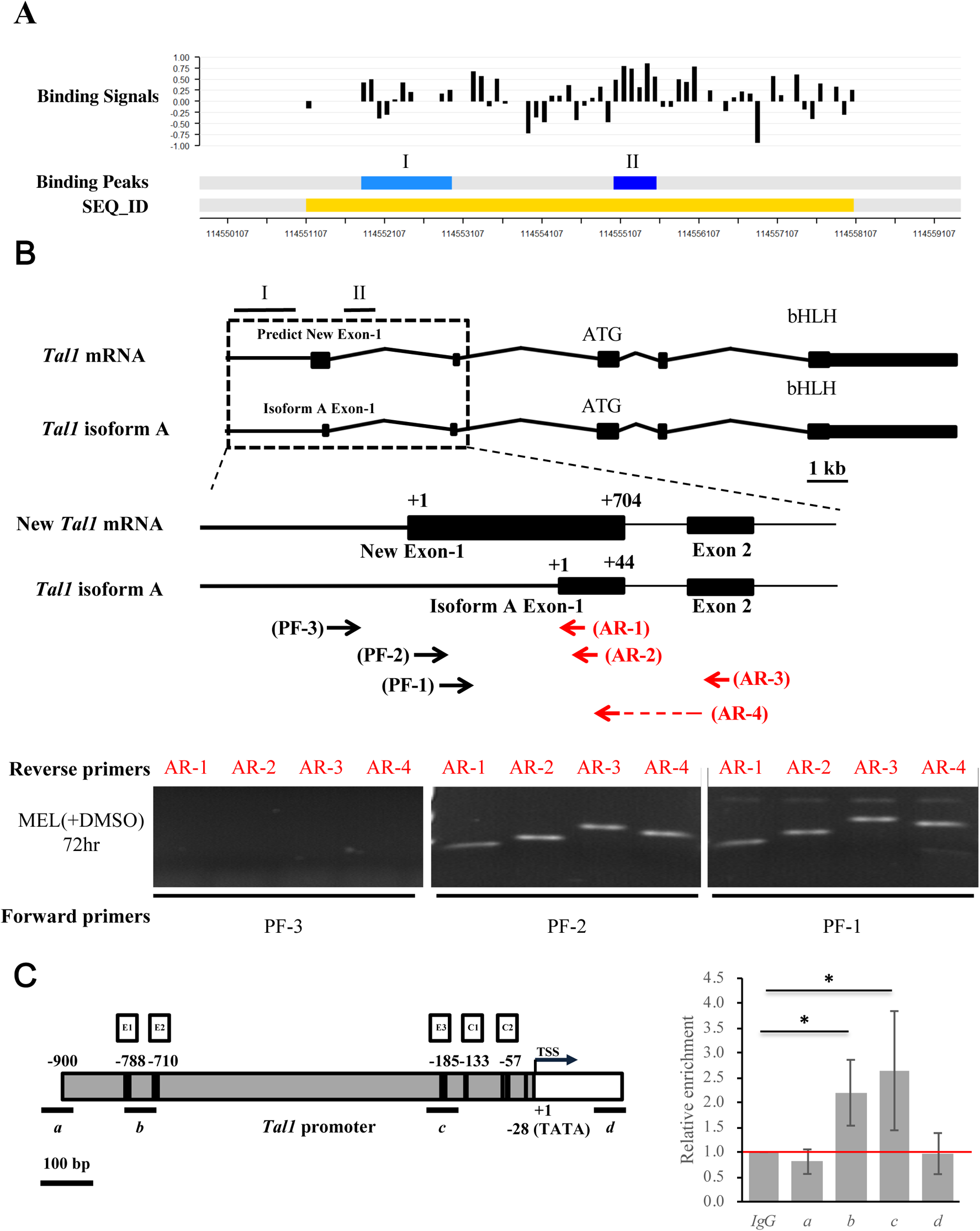
Identification and characterizaton of the authentic exon-1 and promoter of Tal1 gene. **(A)** ChIP-chip promoter array data around the Tal1 gene region. The black bars indicate the signals from the individual probes (“Binding Signals”). The blue regions I and II indicate the EKLF-binding signals from ChIP-chip promoter array analysis of two different mouse E14.5 fetal liver samples (“Binding Peaks”). The yellow region indicates the sequence id (SEQ_ID). **(B)** RT-PCR validation of the newly identified exon 1 of *Tal1* mRNA. Top, maps of the newly identified exon structure of Tal1 mRNA in comparison to that of *Tal1* isoform A. The exons 1-5 are represented by the black boxes. Middle, maps of the exon-1 of *Tal* mRNA and *Tal1* isoform A, respectively, are shown above the primers used for RT-PCR analysis of the *Tal1* mRNA. The sequences of the reverse primers AR-1 and AR-2 are derived from the exon 1 of *Tal1* isoform A. AR-3 is derived from exon 2 sequence. The reverse prime AR-4 is across exons 1 and 2. The sequences of the forward primers PF-1 and PF-2 are from the predicted new exon-1, while PF-3 is from the upstream region. Bottom, gel band patterns of RT-PCR analysis of DMSO-indeed MEL cell RNAs using different sets of PCR primers. **(C)** ChIP-qPCR analysis of EKLF-binding to the *Tal1* promoter. The map of the *Tal1* promoter region is shown on the left, with the CACCC boxes (E1, E2, E3), CCAAT boxes (C1, C2), the TATA box and the transcription start site (TSS, +1) indicated. The 4 regions (a, b, c and d) are bracketed by the 4 primer sets used in qPCR analysis of chromatin from DMSO-induced MEL cells immunoprecipitated with anti-EKLF. The relative folds of enrichment of the chromatin DNA samples pulled down by anti-EKLF are calculated as the Cq values over those derived from use of the IgG and shown in the right histograph. Error bars represent standard deviations from 3~7 biological repeats. The statistical significance of the difference between experimental and control groups was determined by the two-tailed Student *t* test, * p< 0.05.

**Figure 5.**
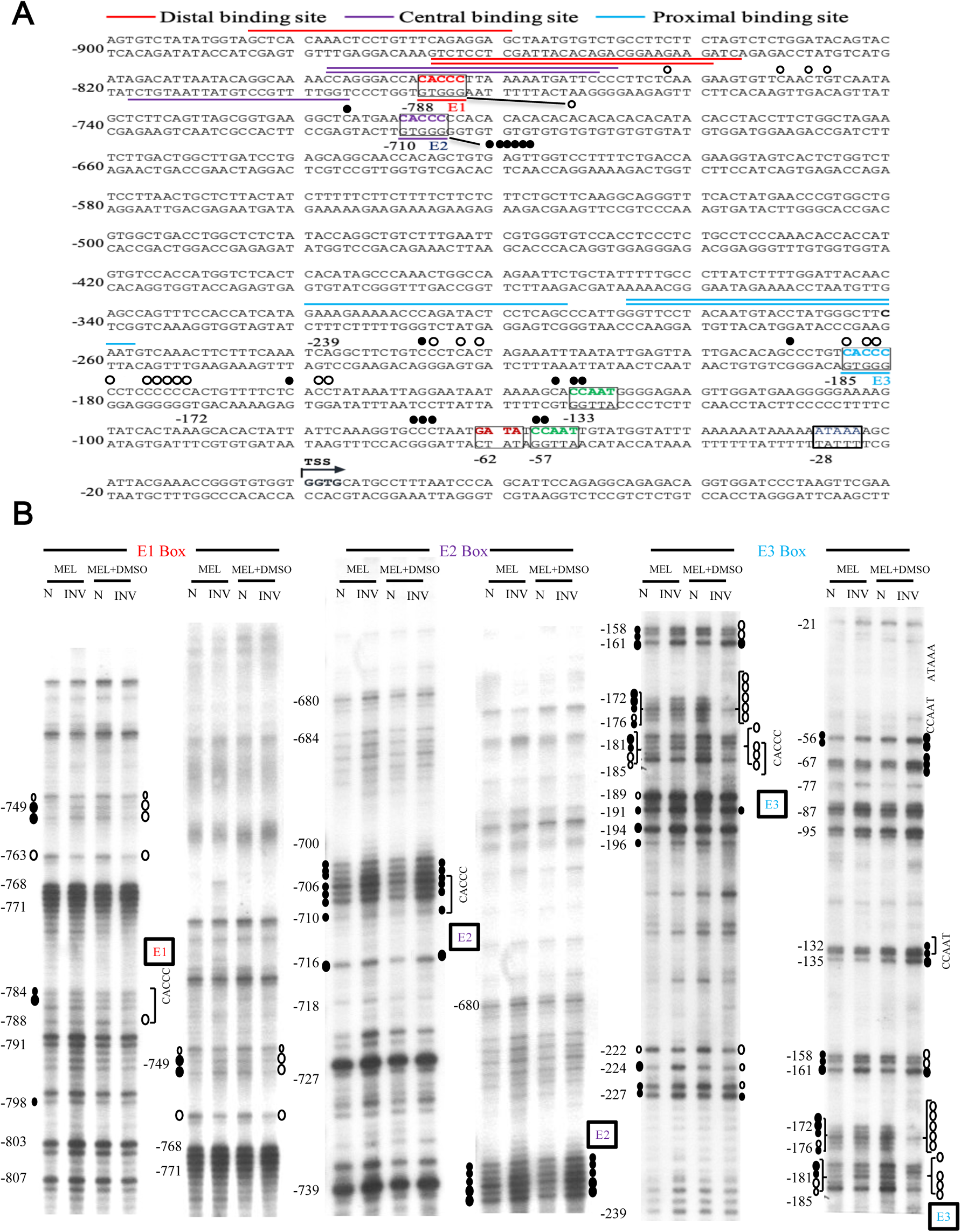
Genomic footprinting analysis of the promoter of Tal1 gene in MEL cells. (A) The protected bases (○) and hyper-reactive bases (●) of the Tal1 promoter region in MEL cells after DMSO induction, as deduced from the in vivo DMS footprinting analysis, are labeled on the DNA sequence. The footprinting pattern indicates the binding of EKLF on the E3 box. (B) The representative autoradiographs of the analysis of the upper strand of the *Tal1* promoter region by *in vivo* DMS protection and LMPCR assay are shown. Locations of different factor-binding motifs/ boxes, i.e., E1, E2, E3, CCAAT, ATAAA, are indicated on the right of the gel patterns. Numbers on the left correlate with those indicated on the sequence in (A). The patterns in the N and INV lanes are the results from *in vitro* and *in vivo* DMS cleavages, respectively. Only those residues consistently showing differences from the controls are indicated. The sizes of the circles reflect the different extents of protection or enhancement of DMS cleavage *in vivo* vs. *in vitro*.

To solve the issue, we carried out semi-quantitative RT-PCR analysis of MEL cell RNAs using different sets of primers. As shown in the bottom panels of Fig. 4B, use of the forward primer PF-3 with any one of 4 different reverse primes (AR-1, AR-2, AR-3, and AR-4) would not generate a RT-PCR band on the gel. On the other hand, the use of the forward primer PF-1 or PF-2 together with the 4 reverse primers generated RT-PCR bands the lengths of which were consistent with the existence of an exon (115,055,766 - 115,056,469, NCBI 36 / mm8) consisting of the previously known isoform A exon 1 at its 3’ region (the diagram, Fig. 4B). Based on these RT-PCR data and the common distance (25-27 bp) between the promoter TATA box and transcription start site(s) of polymerase II-dependent genes, we suggest a map of the promoter region of *Tal1* gene upstream of the newly identified exon1, which contains the TATA box at −28, two CCAAT boxes at −133 and −57, and three CACCC boxes (−788, −710 and −185) upstream of transcription start site or TSS (Figs. 4C and 5A).

To validate the *in vivo* binding of EKLF in the newly identified *Tal1* promoter, we carried out ChIPq-PCR analysis. As shown in Fig. 4C, use of four different sets of primers spanning different regions upstream and downstream of the *Tal1* transcription start site (TSS) indicated EKLF-binding to region b containing the distal CACCC boxes E1 at −788/ E2 at −710 and to region c containing the proximal CACCC box E3 at-185.

### Binding *in vivo* of EKLF to the proximal CACCC box of *Tal1* promoter - Genomic footprinting analysis

In order to examine whether EKLF indeed bound to the proximal CACCC box of the *Tal1* promoter in differentiated erythroid cells, we next carried out genomic footprinting assay of the *Tal1* promoter in MEL cells before and after DMSO induction (Fig. 5). As shown, upon DMSO induction of the MEL cells, genomic footprints appeared at the distal CACCC box E1 and more prominently the proximal CACCC box E3 at-185 (Fig. 5). On the other hand, the distal CACCC box E2 at −710 was not protected in MEL cells with or without DMSO induction. Notably, the intensities of gel bands at −133, −132, −57, and −56 appeared to be enhanced upon DMSO induction, suggesting binding of factor(s) at the two CCAAT boxes as well (Fig. 5). These genomic footprinting data support the scenario that EKLF positively regulates the *Tal1* promoter activity through binding mainly to the proximal promoter CACCC box E3. This would facilitate the recruitment of other factors including the CCAAT box-binding protein(s) to the *Tal1* promoter.

### Requirement of the proximal CACCC motif for transcriptional activation of the *Tal1* promoter by EKLF

To investigate whether EKLF was indeed an activator of *Tal1* gene transcription through binding to the proximal CACCC promoter box, we constructed a reporter plasmid p*Tal1*-luc in which the *Tal1* promoter region from −900 to −1 was cloned upstream of the luciferase reporter. Three mutant reporter plasmids, p*Tal1*(Mut E1)-Luc, p*Tal1*(Mut E2)-Luc, and p*Tal1*(Mut E3)-Luc, were also constructed in which the CACCC box E1, E2, or E3 was mutated (Fig. 6A). Human 293T cells were then co-transfected with one of these 4 reporter plasmids plus an expression plasmid pFlag-EKLF. As shown in Fig. 6B, the luciferase reporter activity in cells co-transfected with p*Tal1*-Luc, p*Tal1*(Mut E1)-Luc or p*Tal1*(Mut E2)-Luc increased in an Flag-EKLF dose-dependent manner. However, mutation at the E3 box of the reporter plasmid p*Tal1*(Mut E3)-Luc prohibited this increase. This result in combination with the genomic footprinting data of Fig. 5 demonstrate explicitly that binding of EKLF to the proximal CACCC box E3, but not the distal E1 or E2 box, in differentiated erythroid cells is required for transcriptional activation of the *Tal1* promoter.

**Figure 6.**
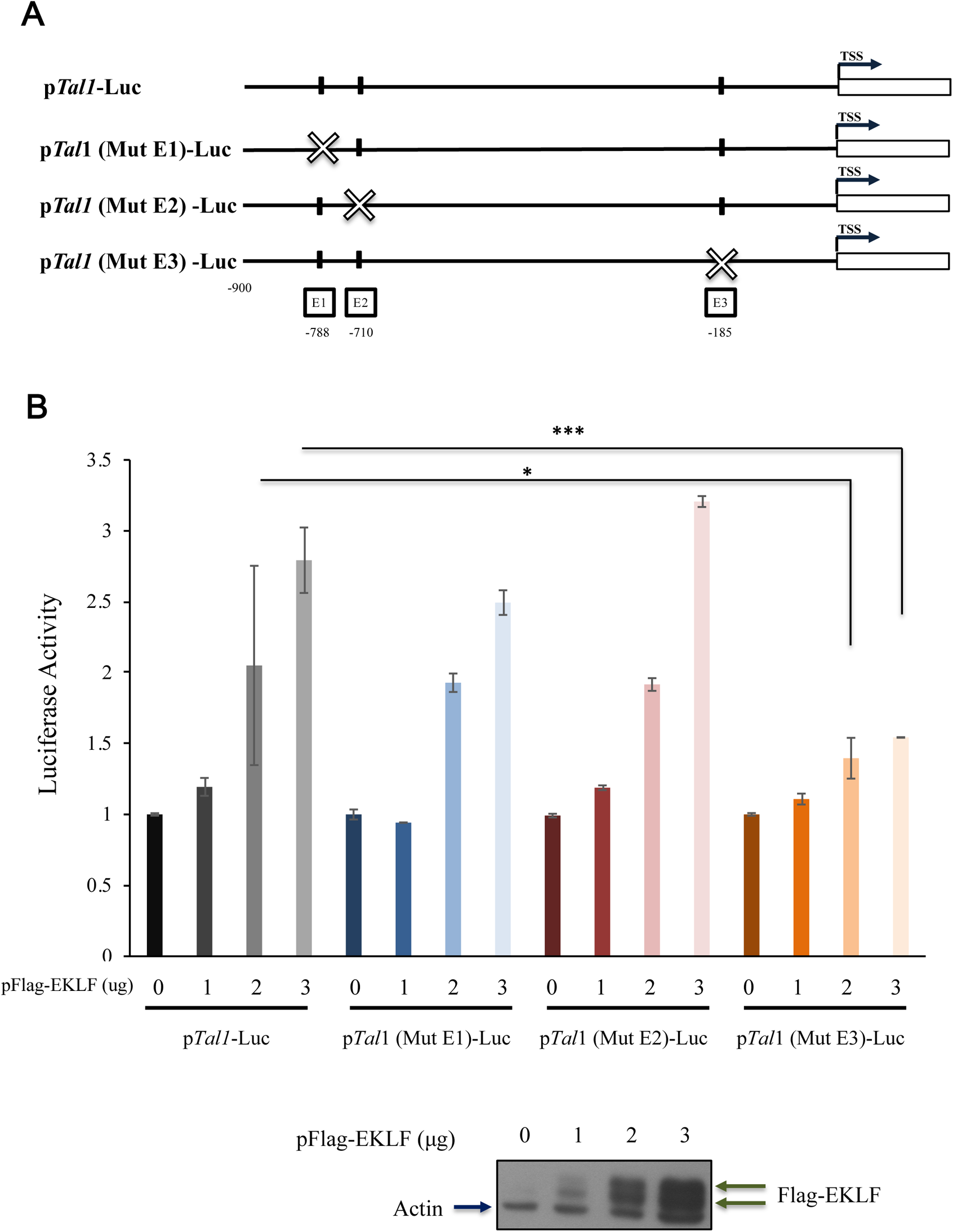
Transactivation of Tal1 promoter by EKLF. **(A)** Linear maps of the wild type and three mutant fragments, in which the CACCC box E1, E2, or E3 was mutated, used for construction of the reporter plasmids p*Tal1*-Luc, p*Tal1*(Mut E1)-Luc, p*Tal1*(Mut E2)-Luc and p*Tal1*(Mut E3)-Luc, respectively. **(B)** Luciferase reporter assay of the *Tal1* promoter in 293 cells. The dose-dependence of the luciferase (Luc) activity on the amount (μg) of pFlag-EKLF used in co-transfection is shown in the bar diagram. *p<0.05, *** p<0.001 by t test. Error bars: STD. The elevated levels of the exogenous Flag-EKLF upon co-transfection with increased amounts of the pFlag-EKLF plasmid were validated by immunoblotting.

## DISCUSSION

A well-coordinated group of transcription factors regulate similar or distinct sets of target genes, which build up the diverse functional networks and biological pathways governing the process of erythropoiesis. Among these factors are GATA1, FOG1, FLI1, PU.1, TAL1/SCL, and EKLF (2–4, 10). Previously, global analyses by gene expression profiling with use of the microarrays have suggested the potential target genes and genetic pathways that function in erythropoiesis, as regulated by GATA1, TAL1/SCL, and EKLF (17, 18, 30, 37, 40, 55). Later, ChIP analysis in combination with next-generation sequencing and microarray hybridization have further provided lists of genes that could be regulated directly, through DNA-binding, by these factors (28, 29, 39, 40,56). Among the factors the potential regulatory targets of which have been studied globally is EKLF. In particular, the two sets of ChIP-Seq analyses have each provided a set of direct target genes of EKLF in the mouse fetal liver cells (28, 29). The change of binding of EKLF to its potential gene targets during differentiation from erythroid progenitors to erythroblasts in the E13.5 fetal liver has also been analyzed (29). However, these two studies have displayed divergent data with respect to the identities of genes directly regulated by EKLF.

n this study, we have analyzed the regulatory functions of EKLF in E14.5 mouse fetal liver cells by combined use of genome-wide expression profiling and promoter ChIP-chip assay. Unexpectedly, the number of direct gene targets (1,866), as defined by the occupancy of EKLF within −3.75kb to +0.75kb relative to TSS (1.2 fold enrichment) and change >1.4 fold of the expression levels upon depletion of *Eklf* in the gene knockout mice, are significantly higher than those derived from Tallack et al, (28) and Pilon et al. (29). As shown in Fig. S7A, of the 1,866 EKLF target genes that we have identified, 257 genes (13.7%) overlap with the data set from Tallack et al. (28) and 231 genes (12.3%) overlap with the data set from Pilon et al. (29). Furthermore, the number of overlapping genes between those two data sets was only 199. Moreover, among the direct targets identified in the 3 studies, only 55 (2.9% of 1,866) are in common (Fig. S7A). The inconsistencies of the conclusions among the 3 groups with respect to the direct target genes of EKLF likely result from the uses of different antibodies, i.e. anti-EKLF from different sources vs. anti-HA, different approaches, ie. microarray hybridization vs. RNA-Seq and ChIP-chip vs. ChIP-Seq., different developmental stages, i.e. E13.5 vs. E14.5, of the embryos analyzed, different mouse strains, i.e. WT vs. HA-EKLF knock-in mice, different cell types, i.e. whole fetal liver vs. the progenitors/erythroblasts, and finally, analysis using different peak calling methods. Moreover, we have used 1.4 fold as the cutoff line, rather than 2 fold chosen by the other two groups (17, 29, 30), when comparing the WT and *Eklf*^−/−^ expression profiles. This lower cutoff line may have allowed us to find more candidate targets that display subtle expression differences but have prominent functional significance. As expected, use of a higher cut-off line, i.e. 2-fold instead of 1.4 fold, for analysis of our ChIP-Chip and microarray data decreased the overlap between our list of direct target genes of EKLF and those derived from the other 2 studies (Supplementary Fig. S7B).

One surprising outcome of our genome-wide study is the existence of a positive feedback loop between the two well-known erythroid-enriched transcription factors, EKLF and TAL1, in early erythroid differentiation. Both factors very likely promote the transition from pro-erythroblasts (Pro-E) to basophilic erythroblasts (Baso-E), as initially suggested by the fact that genetic ablation of either gene in mice causes loss of erythroid cell types beyond the stage of Baso-E (15, 34). Later studies including the use of whole genome ChIP-Seq have further supported *Eklf* being a downstream target of TAL1 (39, 40). By loss-of-function analysis, we show that EKLF also positively regulates the expression of *Tal1* during erythroid differentiation (Fig. 3). In particular, induced depletion of *Eklf* drastically lowers the expression level of *Tal1* in DMSO-induced MEL cells (Fig 3C). The combined data from the ChIP-chip, genomic footprinting, and transient reporter assays further indicate that EKLF activates *Tal1* gene transcription through binding to the proximal CACCC box in the newly identified *Tal1* promoter (Figs. 4 and 5). Consistent with this scenario of mutual activations of *Tal1* and *Eklf*, the mRNAs of *Tal1* and *Eklf* are both progressively up-regulated during erythroid differentiation of the primary mouse fetal liver cells (57). Thus, our finding of the positive regulation of *Tal1* gene by EKLF demonstrates the existence of a *Tal1*-*Eklf* positive feedback loop that promotes the mammalian erythroid differentiation in a tightly regulated time window, from the transition of Pro-E to Baso-E of the erythroid lineage.

We propose the following scenario for the mutual activation of *Tal1* and *Eklf*, and the functional consequences of this positive feedback loop during erythroid differentiation. TAL1 is a predominant transcriptional factor responsible for the origin of the definitive hematopoietic stem cells as well as the differentiation of the erythroid/ megakaryocytic lineages. Its expression pattern spans the HSC cells, the subsequent multipotent progenitors, as well as the erythroid/ megakaryocytic lineages (58). On the other hand, EKLF is expressed in the erythroid cells, megakaryotes, hematopoietic stem cells (HSC), as well as in hematoprogenitors including MEP, GMP, and CMP (6, 59, Bio GPS). In the erythroid lineage at the BFU-E/CFU-E/Pro-E stages, the two factors are already expressed at basal levels. EKLF is retained by FOE in the cytosol (26), while TAL1 protein positively regulates the expression of *Eklf*, as suggested by the previous reporter assay (41) and ChIP-Seq data of TAL1 binding on the *Eklf* promoter in BFU-E/CFU-E/Pro-E (39, 40). When the cells enter the Baso-E stage, EKLF protein is released from its physical interaction with FOE in the cytoplasm and imported into the nucleus (26). The imported EKLF binds to the E3 box of the *Tal1* promoter to enhance the promoter activity of *Tal1* (Fig. 5 and 6). This positive feedback loop would rapidly amplify both factors during erythroid terminal differentiation. As a result, the EKLF-mediated activation of *Tal1* may act as a valve that facilitates the commitment of the erythroid lineage from MEP through promoting the differentiation transition from Pro-E to Baso-E, thus sustaining the process after Baso-E. Furthermore, there is a high frequency of co-occupancy of EKLF and TAL1 in a number of promoters that are active in erythroid cell lines or erythroid tissues (Table 4; 28–30). Thus, the *Eklf*/*Tal1* loop would irreversibly promote erythroid terminal differentiation through the up-regulation of not only *Eklf* and *Tal1*, but also their mutual downstream targets that are crucial for erythroid differentiation, such as *Hba, Hbb, E2f2*, *Gypa, Epb4.1*,and *Alas2*, among others. For the latter process, the EKLF and TAL1 proteins may work within the same transcriptional complex(es) which binds to the composite CACCC box-E box in the promoters of these downstream targets (51).

In sum, positive feedback loops of transcriptional regulation are essential for the progression of different physiological processes, as shown previously for *c-kit* with *Tal1* in the survival and clonal expansion of the progenitor hematopoietic cells (60), early B cell factor (EBF) with MyoD in the commitment and differentiation of *Xenopus* muscle cells (61), and *c-Myc* with *Sox2* in the self-renewal of mouse multipotent otic progenitor cells (62), etc. In comparison to the above cases, the cooperation between EKLF and TAL1 in the promotion of erythroid differentiation provides an unique case for ensuring the commitment to erythroid differentiation among the multiple lineages of the hematopoietic system.

## ACKNOWLEDGEMENTS

We thank our laboratory colleagues Yu-Chi Chou, Keh-Yang Wang and An-Chun Lee for providing us various reagents and advices. The generosity by Dr. François Morlé at Université de Lyon, Lyon, France in sharing the stable MEL cell clones 4D7/2M12 and the experimental procedures is deeply appreciated. We also thank Dr. Shu-Yun Tung at Genomics Core Facility, Institute of Molecular Biology, Academia Sinica for her help with the analysis of Mouse Genome Array 430A 2.0 (Affymetrix, Inc.). This research was supported by grants from the National Health Research Institute and the Academia Sinica (AS), Taipei, Taiwan, Republic of China. The works on bioinformatics were supported by Mathematics in Biology Group of Institute of Statistical Science, AS and by grants from Academia Sinica, AS-100-TP-AB & AS-104-TP-A07. C.-K. James Shen was an NSC Frontier of Science Awardee and AS senior investigator Awardee.

## AUTHORSHIP CONTRIBUTIONS

C.-H. Hung, Y.-S. Huang, T.-L, Lee, and K.-C. Yang contributed equally to this work; Y.-S. Huang and J.-H. Hung designed and performed the experiments and analyzed the data regarding to the regulatory relationship between *Eklf* and *Tal1*; T.-L. Lee, K.-C. Yang, and S. Yuan performed the whole-genome analyses of datasets obtained from NimbleGen ChIP-chip assay and Affymetrix gene expression profiling. S.-C. Wen performed the genomic footprinting analysis. J.-H. Hung, Y.-C. Shyu, M.-J. Lu, and S.-C. Wen carried out the validation experiments of the data from whole-genome analyses. Y.-S. Huang, J.-H. Hung, T.-L. Lee, S. Yuan, and C.-K. J. Shen developed the project and wrote the manuscript.

## FIGURE LEGENDS-S

**Figure S1. <u>Associated functional networks of up-regulated EKLF targets in E14.5 fetal liver cells.</u>**

IPA was performed on the data from comparing EKLF promoter ChIP-chip and microarray data from wild-type and *Eklf^−/−^* E14.5 fetal liver cells, using 710 up-regulated EKLF target genes as the focus gene set. The highest-scoring network was **(A)** lipid metabolism, molecular transport, small molecule biochemistry and cellular development, cellular growth and proliferation with the significance score 34. The secondary network was **(B)** hematological system development and function were eligible (score 26). Arrows and lines denote interactions between specific genes within the network. Solid lines indicate direct relationships, and dashed lines indicate indirect relationships.

**Figure S2. <u>Associated functional networks of down-regulated EKLF targets in E14.5 fetal liver cells.</u>**

IPA was performed on the data from comparing EKLF promoter ChIP-chip and microarray data from wild-type and *Eklf^−/−^* E14.5 fetal liver cells, using 1,156 down-regulated EKLF target genes as the focus gene set. The highest-scoring network was (A) carbohydrate metabolism, lipid metabolism, small molecule biochemistry and (B) cancer, tumor morphology, post-translational modification both have score 34. Other associated network functions include (C) developmental disorder, hereditary disorder, immunological disease; (D) endocrine system development and function, molecular transport, protein synthesis; and (E) organismal injury and abnormalities, skeletal and muscular disorders, cell death and survival. Arrows and lines denote interactions between specific genes within the network. Solid lines indicate direct relationships, and dashed lines indicate indirect relationships.

**Figure S3. <u>RT-PCR validation of mouse fetal liver gene expression in KO embryos vs. WT embryos.</u>**

Bar diagram of the relative mRNA levels of each candidate genes in E14.5 fetal liver cells of the *Eklf^−/−^* (KO) mice and WT mice, as analyzed by semi-RT-PCR and normalized by *actin*. * p < 0.05, ** p < 0.01, *** p < 0.001 by t test. Error bars, S.E.M.

**Figure S4. <u>Validation of ChIP-chip data by ChIP-qPCR.</u>**

ChIP-Q-PCR analysis of EKLF-binding promoter regions from ChIP-chip binding peaks. The statistical analysis of the ChIP-Q-PCR data is shown in the bar diagrams. The promoter region signals were compared with IgG control. * p < 0.05, ** p < 0.01, *** p < 0.001 by t test. Error bars, S.D.

**Figure S5. <u>IPA identified significant canonical pathways associated with the up-regulated EKLF targets in E14.5 fetal liver cells.</u>**

The five most significant canonical pathways obtained by IPA were (A) cell cycle control of chromosomal replication; (B) EIF2 signaling; (C) mitochondrial dysfunction,; (D) hypusine biosynthesis, and (E) tryptophan degradation III (Eukaryotic). The upregulated EKLF targets are indicated in yellow.

**Figure S6. <u>IPA identified significant canonical pathways associated with the down-regulated EKLF targets in E14.5 fetal liver cells.</u>**

The five most significant canonical pathways obtained by IPA were (A) insulin receptor signaling; (B) chondroitin sulfate degradation (Metazoa); (C) gap junction signaling; (D) Nitric Oxide signaling in the cardiovascular system, and (E) PDGF signaling. The down-regulated EKLF targets are indicated in yellow.

**Figure S7. <u>Three-way Venn diagrams displaying the overlap of direct targets of EKLF as derived from Tallack et al. (28), Pilon et al. (29), and the current study.</u>**

(A) Diagram with cut-off line of 1.4 fold

(B) Diagram with cut-off line of 2.0 fold

## LIST of SUPPLEMENTARY TABLES

**Supplementary Table 1.** List of EKLF-bound regions on mouse genome.

**Supplementary Table 2A.** List of EKLF direct downstream targets generated from matching between the Affymetrix expression profiling comparing wild-type and *Eklf^−/−^* (KO) mice and the NimbleGen ChIP-chip promoter array.

**Supplementary Table 2B.** List of EKLF up-regulated genes after filtering with effect sizes.

**Supplementary Table 2C.** List of EKLF down-regulated genes after filtering with effect sizes.

**Supplementary Table 3A.** List of top enriched network functions from up-regulated genes derived from Ingenuity IPA software.

**Supplementary Table 3B.** List of top enriched network functions from down-regulated genes derived from Ingenuity IPA software.

**Supplementary Table 4A.** List of molecular and cellular functions from up-regulated genes derived from Ingenuity IPA software.

**Supplementary Table 4B.** List of molecular and cellular functions from down-regulated genes derived from Ingenuity IPA software.

**Supplementary Table 5A.** List of significant canonical pathways from up-regulated genes derived from Ingenuity IPA software.

**Supplementary Table 5B.** List of significant canonical pathways from down-regulated genes derived from Ingenuity IPA software.

**Supplementary Table 6.** The occurrences of binding motifs of 226 transcription factors in EKLF-bound regions.

**Supplementary Table 7.** List of primers used in ChIP-chip data validation by ChIP-qPCR.

**Supplementary Table 8.** List of RT-PCR primers of used for validation of microarray hybridization data.

**Supplementary Table 9.** List of primers used for identification of Tal1 exon-1.

